# Detecting hierarchical 3-D genome domain reconfiguration with network modularity

**DOI:** 10.1101/089011

**Authors:** Heidi K. Norton, Harvey Huang, Daniel J. Emerson, Jesi Kim, Shi Gu, Danielle S. Bassett, Jennifer E. Phillips-Cremins

## Abstract

Mammalian genomes are folded in a hierarchy of topologically associating domains (TADs), subTADs and looping interactions. The nested nature of chromatin domains has rendered it challenging to identify a sensitive and specific metric for detecting subTADs and quantifying their dynamic reconfiguration across cellular states. Here, we apply graph theoretic principles to quantify hierarchical folding patterns in high-resolution chromatin topology maps. We discover that TADs can be accurately detected using a Louvain-like locally greedy algorithm to maximize network modularity. By varying a resolution parameter in the modularity quality function, we accurately partition the mouse genome across length scales into a hierarchical nested structure of network communities exhibiting a wide range of sizes. To distinguish high probability subTADs from the full detected set, we developed and applied a new ‘hierarchical spatial variance minimization’ method. Moreover, we identified a large number of dynamically altered communities between pluripotent embryonic stem cells and multipotent neural progenitor cells. Cell type specific boundaries correlate with trends in dynamic occupancy of the architectural protein CTCF, thereby validating their biological relevance. Together, these data demonstrate the utility of metrics from network science in quantifying a nested hierarchy of dynamic 3D chromatin communities across length scales. Our findings are significant toward unraveling the link between higher-order genome folding and gene expression during healthy development and the deregulation of molecular pathways linked to disease.

## Introduction

Principles from a field of mathematics known as graph theory have emerged as powerful tools for quantifying connectivity patterns within complex systems (Bollobás 1985). Networks are graphs consisting of nodes connected by edges that can be used to represent the underlying structure of biological, social, physical and information systems (Newman 2010). Complex networks often show hierarchical patterns of connectivity across length scales (Ravasz and Barabási 2003). A node can connect to an interacting neighbor via a direct link. Neighboring nodes can form small subnetworks arranged in distinct recurrent configurations termed motifs (Sporns and Kötter 2004). Subnetworks can in turn aggregate into modular structures termed communities in which the nodes in a particular community are more connected to each other than expected by chance (Porter, Onnela, and Mucha 2009; Lancichinetti and Fortunato 2009). Small-scale communities can further aggregate into larger-scale communities to create nested, hierarchical structures (Bassett et al. 2010). Network science offers tools to identify such structures, and has therefore been widely employed to characterize the connectivity patterns of real-world complex systems across length scales.

A complex system exhibiting hierarchical structure in the field of biology is the 3-D folding of the genetic material. In mammalian nuclei, the ~2-meter-long genome is arranged into complex higher-order configurations to fit inside a nucleus that is 5-10 *μ*m in diameter. Seminal studies based on microscopy (Fraser and Bickmore 2007; Kosak and Groudine 2004; Lanctôt et al. 2007) or molecular proximity ligation and deep sequencing (Dostie et al. 2006; Lieberman-Aiden et al. 2009; Dixon et al. 2012; Simonis et al. 2006; Fullwood et al. 2009) have revealed that chromatin folding is nonrandom with unique patterns across disparate length scales. At the largest length scale, chromosomes fold into independent territories with respect to each other (Cremer and Cremer 2001). Active and inactive genomic regions then partition into large-scale A/B compartments distinguished by transcriptional activity and activating or repressing chromatin modifications (Lieberman-Aiden et al. 2009). Within compartments, chromatin is folded into topologically associating domains (TADs) in which genomic loci within ~1 Megabase (Mb)-sized regions contact each other more frequently than neighboring regions to form globular, domain-like structures (Dixon et al. 2012; Nora et al. 2012). Within TADs, smaller folding domains termed subTADs have been uncovered (Phillips-Cremins et al. 2013; Rao et al. 2014). Moreover, looping interactions, in which distal genomic segments are spatially connected and loop out the intervening DNA, occur within and between subTADs (Rao et al. 2014; Dowen et al. 2014; Morey et al. 2007). At the smallest length scale, DNA wraps around the histone octamer to form the 10 nm chromatin fiber. How the nested, hierarchical nature of chromatin architecture is linked to genome function remains an important unanswered question with significant implications for understanding the molecular mechanisms governing development and disease.

Genome folding at the sub-Mb scale within TADs remains poorly understood, due in part to the low resolution of proximity ligation maps and the paucity of methods to robustly quantify subTAD domain organization. Here, we quantify the hierarchically modular structure of the 3-D genome by combining (i) high resolution (~4 kb) genome folding maps, (ii) mathematical principles from network science and (iii) cellular states representing two different stages of neural development. We conceptualize proximity ligation data as a network with each 4 kb genomic bin represented as a node and each interaction frequency between two nodes represented as an edge. By employing a community detection method based on the optimization of the network modularity metric (Newman 2006), we find that the nodes cluster into a nested hierarchy of communities. Many smaller sub-Megabase-sized communities restructure during the switch from pluripotent Embryonic Stem (ES) cells to multipotent primary neural progenitor cells (pNPCs), whereas some communities are invariant between the two cellular states. Dynamic community boundaries correlate with dynamic occupancy for a known architectural protein, thus supporting the validity of our modularity optimization algorithm for detecting genomic domains. The application of network science to genome topology data offers an orthogonal approach to traditional proximity ligation analysis methods that could enhance discovery of biologically relevant differences in genome organization in cellular models of development and disease.

## Results

### 3D genome folding data as a network

Social (Zachary 1977; Traud, Mucha, and Porter 2011), technological (Balthrop et al. 2004) and biological (Greicius et al. 2003; Segal et al. 2003) systems have been modeled as complex networks of nodes interconnected by edges weighted by the strength of node-node interactions. Because genome-wide proximity ligation data is also inherently relational data, we reasoned that network mathematics might have utility in capturing key features of higher-order chromatin folding. To test this idea, we abstracted the genome’s higher-order configurations as a square, symmetric adjacency matrix, *A*, with the nodes representing adjacent genomic loci parsed into equally-spaced bins. *A* is an *NxN* matrix where *N* is the total number of nodes. Edges represent the frequency of interaction, *A_ij_*, between node *i* and node *j* across a population of cells (**Figure 1**). This representation of genome topology as a network opens up possibilities for inquiry via a range of graph theoretical tools.

**Figure 1.**
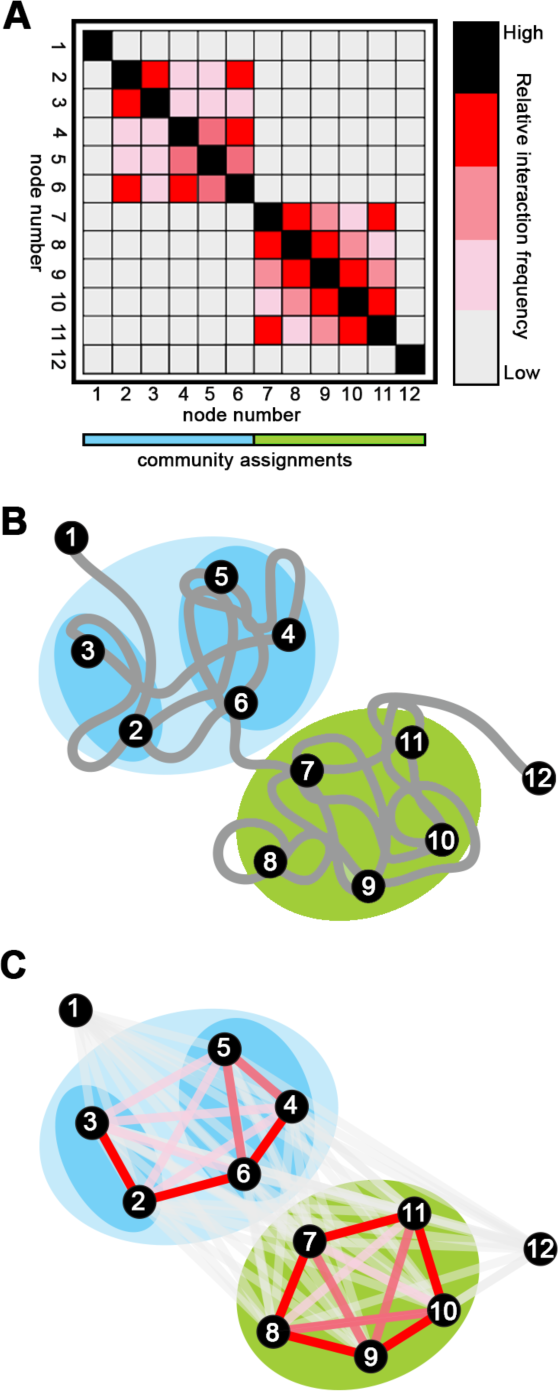
3D genome folding data can be represented mathematically as a network. (A) Schematic representation of a HiC or 5C heatmap with two Mb-scale topologically associated domains (TADs) and nested sub-TADs within the first TAD. Node numbers correspond to DNA restriction fragments. (B) Model of the potential community structure underlying proximity ligation data depicted in panel (A). Black dots with numbers represent nodes corresponding to the genomic coordinates represented in each bin. (C) Spring force diagram representation of the network. Edge colors depict edge weight ranging from low (grey) to high (red).

### Community detection in 3-D chromatin folding networks

We first set out to understand the general network structure of genome architecture data by visualizing heatmaps of low-resolution (40 kb binned) Hi-C data generated from mouse cortex (Dixon et al, 2012). Hi-C is a genome-wide molecular assay that relies on proximity ligation and high-throughput sequencing to detect discrete counts representing the frequency with which two genomic segments interact across a population of cells (Dekker et al. 2002; de Wit and de Laat 2012; Lieberman-Aiden et al. 2009). Visual inspection of matrices representing 60 Mb-sized regions around the *Sox2* and *Olig1/Olig2* genes confirmed folding patterns consistent with previously reported TADs and nested subTADs (Dixon et al. 2012; Nora et al. 2012; Phillips-Cremins et al. 2013; Sanborn et al. 2015) (**Figure 2A**). We next overlaid published TADs called with a directionality index metric and a Hidden Markov Model on the heatmaps (**Figure 2A**, blue lines, Dixon et al. 2012).

**Figure 2.**
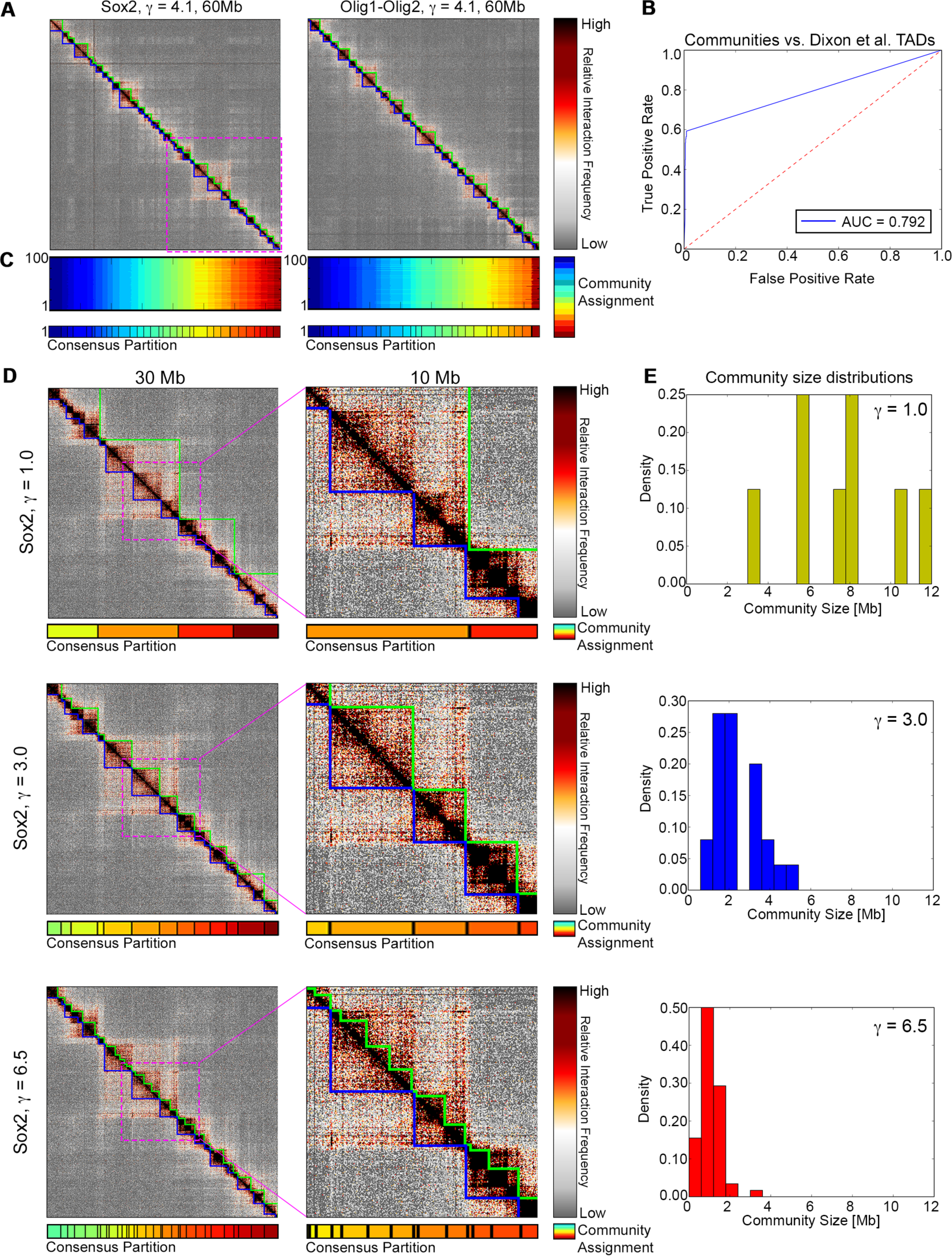
Network modularity maximization identifies TADs. (A) Hi-C heatmaps of mouse cortex from Dixon et al. 2012 in 60 Mb regions surrounding Sox2 (left) and Olig1-Olig2 (right) genes. TAD calls reported by Dixon et al. 2012 are outlined in blue on the heatmap. Communities identified by applying the Louvain-like locally greedy algorithm to maximize modularity with structural resolution parameter *γ* = 4.1 one hundred times and determining a consensus partition are outlined in green. (B) Receiver operating characteristic (ROC) curve to assess the performance of network modularity maximization in calling TADs previously published by Dixon et al. 2012. Communities identified with network modularity maximization in the 60 Mb Sox2 and 60 Mb Olig1-Olig2 regions were pooled together and compared to Dixon et al. TAD calls within the same two 60 Mb regions. The area under the curve (AUC) is 0.792. (C) One hundred underlying partitions (top) used to determine the consensus partition (bottom) for each genomic region. (D) Hi-C heatmap of a 30 Mb region (left) and 10 Mb region (right) surrounding the *Sox2* gene with consensus partitions determined at three different resolution parameter values: *γ* = 1.0, 3.0, and 6.5 plotted beneath the heatmaps. Green lines on the heatmap indicate the communities identified from each consensus partition. Similar to panel (A), TAD calls from Dixon et al. 2012 are outlined in blue. (E) Size distributions of the communities called in the 60 Mb Sox2 region for the three different resolution parameter values shown in panel (D).

We noticed that the Dixon et al. TADs showed strong qualitative concordance with intermediate Mb-sized domains, whereas larger Megadomains and smaller/nested subTADs remained unclassified. Several subsequent methods for domain calling have been reported, achieving similar performance in calling Mb-sized TADs to the seminal Dixon et al. approach (Weinreb and Raphael 2016; Filippova et al. 2014; Lévy-Leduc et al. 2014; Rao et al. 2014). However, to date, there is a paucity of methods for (1) detecting chromatin domains of varying sizes assembled in a nested, hierarchical manner without *a priori* knowledge of the number or size of communities, and (2) accurately quantifying dynamic community structure across biological conditions and perturbations.

Networks can be decomposed into groups of highly interconnected subnetworks called communities or modules. Communities are important conceptually because they might represent the grouping of seemingly disconnected nodes into higher-level functional units. The problem of module detection and subsequent thresholding into *bona fide* communities represents a challenging and active area of research, for which there are numerous algorithmic solutions. One of the most widely adopted methods for community detection in the field of network science involves the iterative maximization of a metric of community strength in order to identify the ‘optimal’ partition of nodes into communities (Blondel et al. 2008). Here, to identify communities in our 3D genome folding data, we maximize the modularity quality index, *Q* (1):

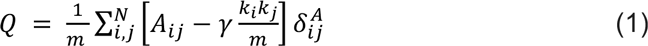

where *A_ij_* is the edge weight representing interaction frequency between nodes *i* and *j*, *k_i_* is the sum of all edge weights for node *i*, *m* is the total sum of all edge weights in the network ***A*** (i.e., the sum of all interaction counts in the matrix), *N* is the total number of nodes in matrix *A* and *γ* is a resolution parameter (discussed below). To ensure that only edges within communities are added to the summation, the Kronecker delta, 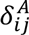, is 1 if *i* and *j* are assigned to the same community and 0 otherwise. The intuition for the modularity metric is that it represents the proportion of edges that fall within communities above that expected in an appropriate null model (M. E. J. Newman and Girvan 2004). A modularity value close to 1 indicates strong community structure and a high quality division of the network into communities, whereas a modularity value close to 0 indicates that the strength of within-community connections is no higher than would be expected by chance.

To identify the optimal division (or “partition”) of the network into communities, we employed a Louvain-like, locally greedy modularity maximization algorithm (Blondel et al. 2008). This algorithm was chosen because it does not require *a priori* knowledge of the number of communities that will be detected (M. E. J. Newman 2011) and it offers the critical capability of identifying nested communities through the variation of a structural resolution parameter (Lancichinetti and Fortunato 2009; Reichardt and Bornholdt 2006) (detailed in **Materials and Methods)**. Briefly, the algorithm begins at iteration t = 0 by assigning each node to its own community. The modularity value associated with this initial partition is computed using Equation (1). Nodes are then merged into communities by randomly selecting a node and merging it with the neighboring node that leads to the greatest gain in modularity (if any). Once every node has been given the opportunity to form a community by merging with another node, the iteration is complete. At the end of the iteration, the gain in modularity, ΔQ, is computed as (2):

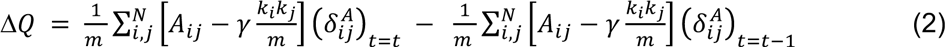

where *t* is the iteration number. If *ΔQ* exceeds a threshold of 1*x*10^−10^, the algorithm continues to the next iteration in which communities are merged with other communities. The iterations continue until no single merge leads to a gain in modularity that exceeds the threshold, at which point the resulting partition of nodes into communities is an ‘optimal’ community partition. In this optimal partition, nodes have been grouped into a set of communities that results in a local or global maximum of the modularity metric.

### Network communities across length scales in low resolution HiC data

We first set out to assess the performance of the modularity optimization method in identifying TADs in 40 kb Hi-C data (Dixon et al. 2012) (**Figure 2A, B**). We observed that the community boundaries closely mirrored the gold-standard Dixon et al. TAD calls (**Figure 2A**). The community partitions were both sensitive and specific, as assessed by the Receiver Operator Characteristic Area Under the Curve (ROC-AUC) of 79% (**Figure 2B**). Moreover, we found that in nearly every case of disagreement between our method and the gold standard Dixon et al. method, the communities identified across the two methods formed nested structures within each other. (**Supplementary Figure 1**). These data provided the initial indication that modularity optimization might have utility in identifying domain structures in proximity ligation data.

A known limitation of the modularity optimization approach is the possibility of converging on a locally maximal modularity value, leading to degeneracies in identifying ‘optimal’ community partitions (Blondel et al 2008; Lancichinetti et al 2009; Good, de Montjoye, and Clauset 2010). Indeed, we noticed variation in the community partition emerging from our modularity optimization algorithm due to random selection of the starting node at the beginning of each run. To address the boundary variation due to convergence on different local maxima, we sampled the landscape of community partition solutions by applying our Louvain-like, locally greedy algorithm 100 times (hereafter referred to as a ‘partition block’ with dimensions 100 x N). We then identified a “consensus partition” by (i) computing a similarity score (the Adjusted Rand Index (aRAND)) between all pairs of partitions within a partition block and (ii) selecting the partition with the maximal average aRAND (Doron, Bassett, and Gazzaniga 2012, detailed in **Materials and Methods**). We observed that the final community assignments from the consensus partition largely agreed with the underlying partition block (**Figure 2C**). Thus, we can address fluctuations in partitions due to algorithmic convergence on local vs. global modularity maxima by computing a “consensus community partition”.

We next hypothesized that we might identify genome communities of different sizes through modification of the structural resolution parameter *γ* (Reichardt and Bornholdt 2006). Intuitively, when *γ* = 1, the positive contribution of intra-community edges are weighted equally to the negative contribution of the null model’s expected edge weight (**Equation 1**). When *γ* > 1, the negative contribution of the expected edge weight is amplified in the modularity equation, thus resulting in lower modularity values for a given partition. Upon modularity maximization the potential gain in modularity for a single move is also lower, making it more difficult to satisfy the *ΔQ* threshold. The resulting ‘optimal’ partition will in turn be biased towards smaller communities. Conversely, when *γ* < 1, it becomes easier to satisfy the *ΔQ* and modularity maximization is biased towards detection of larger communities. We explored a range of *γ* values and noticed that our modularity optimization algorithm could identify large, medium and small domain-like structures in a manner that was inversely proportional to the magnitude of *γ* (**Figure 2D**). Indeed, at *γ* values of 1.0, 3.0, and 6.5, the median community size was 7660, 2160 and 980 kb, respectively (**Figure 2E**). Notably, at low *γ* values, it appeared that the method captured the higher-order level of genome folding termed ‘A/B compartments’, thus further highlighting the importance of optimizing *γ* to the level in the genome folding hierarchy under investigation. These results suggest that the modularity maximization method can accurately detect chromatin folding domain structures in a nested hierarchy across genomic length scales through the simple modification of only a single structural resolution parameter.

### Identification of overlapping subTADs in high resolution 5C data

To date, the accurate identification of subTADs has been challenging, in part due to the complexities of the overlapping and nested domain structure at the subMb-scale. We reasoned that the structural resolution parameter would enable us to explore the complexities of finer scale genome folding at the sub-Mb scale within TADs. To test this idea, we used high-resolution (4 kb) interaction frequency matrices generated by 3C variant termed Chromosome-Conformation-Capture-Carbon-Copy (5C) (Beagan et al. 2016). 5C involves a hybrid capture step in which the ligation junctions from specific genomic regions of interest are selected and amplified from a larger population of all genome-wide ligation junctions (Dostie et al. 2006; Nora et al. 2012; J. E. Phillips-Cremins et al. 2013). The key advantage of 5C technology is the ability to map higher-order chromatin interactions at 4 kb resolution with a fraction of the reads needed for genome-wide Hi-C. For example, the 4 kb-binned 5C map in primary neural progenitor cells spanning a 1 Mb-sized region around the critical developmentally regulated genes *Olig1* and *Olig2* reveals complex fine scale genome folding patterns and clear subTAD structures nested within the larger TAD (**Figure 3A**).

**Figure 3.**
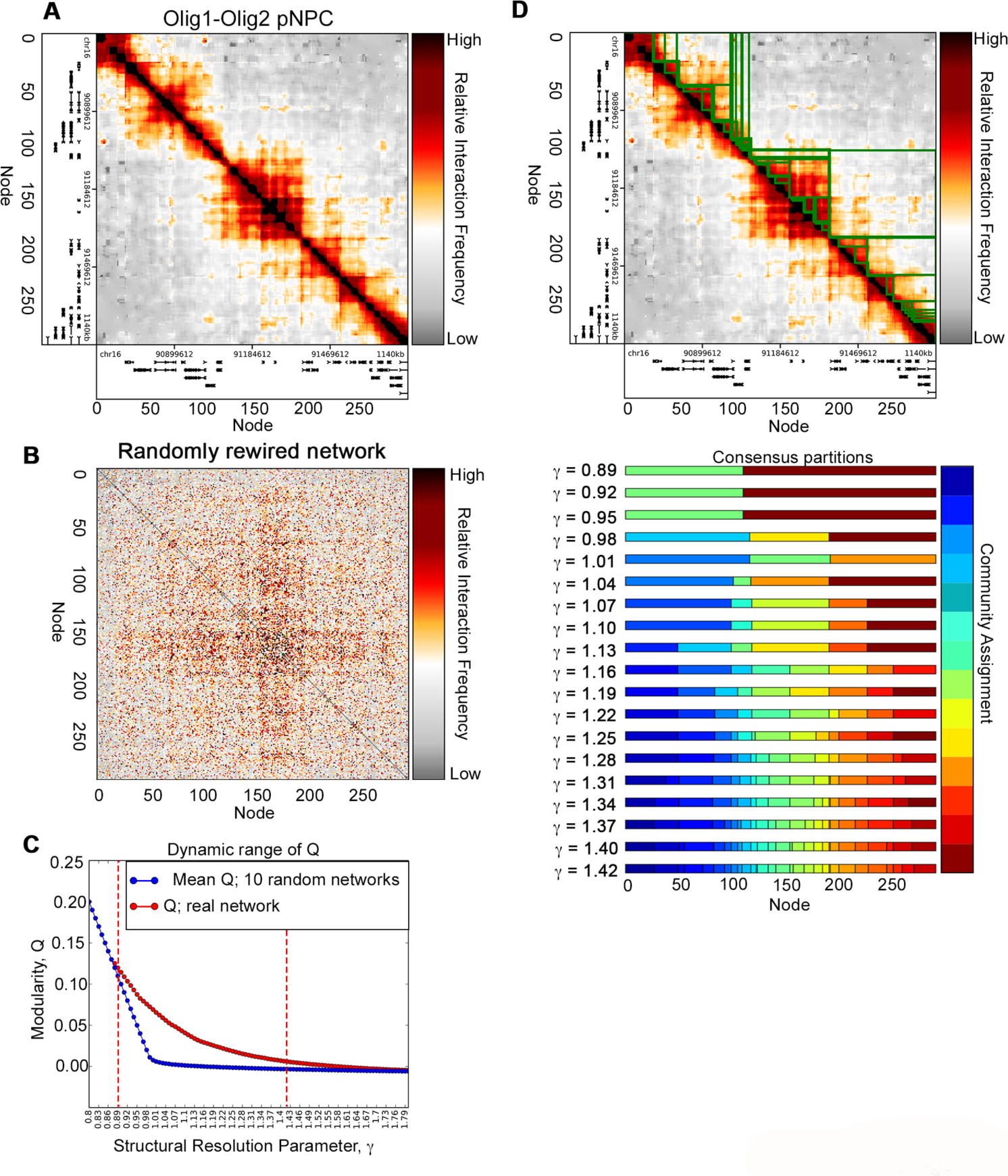
Variation of structural resolution parameter, *γ*, partitions 5C genome folding networks into a nested hierarchy of sub-Mb communities. (A) 5C heatmap of a ~1Mb region surrounding Olig1-Olig2 in primary mouse neural progenitor cells. Genes in the region are plotted on the x and y axes. Network coordinates in terms of genomic coordinates and node number are shown beneath the heatmap. (B) Heatmap of the network from panel (A) that has been randomly rewired by reassigning the location of existing edges according to expected edge weights. (C) Modularity, *Q*, versus the resolution parameter, *γ*, value for input and randomly rewired networks. Mean modularity of 10 randomly rewired networks at each value of the structural resolution parameter, *γ*, is shown in blue, whereas the modularity of the input network from panel (A) is shown in red. Red dashed lines indicate the dynamic range of *γ* values for which the difference between the modularity of the real network and the modularity of the random networks is the largest (**see Supplementary Figure 2**): 0.89 − 1.42. (D) Consensus partitions of Olig1-Olig2 into communities across the *γ* range, 0.89 − 1.42. Community assignments indicated by different colors in the consensus partitions are outlined on the heatmap in green.

To determine if our modularity optimization method has utility in identifying small-scale community structure within TADs, we began by comparing the network properties of real and randomly rewired networks. We generated randomly rewired undirected, weighted graphs with preserved weight, degree, and strength distributions via an established rewiring algorithm (Maslov and Sneppen 2002; Rubinov and Sporns 2011) (**Figure 3B**). For each 5C region, we determined the useful range of *γ* values by comparing the modularity metric of real and randomly rewired networks (**Figure 3C, Supplemental Figure 2**). In the *Olig1/Olig2* network in pNPC cells, *Q* for the real network diverges from the random network at *γ* = 0.88 and converges within 15% of the random network at *γ* = 1.43. On the basis of these properties, we selected the range of *γ* values for community detection at *Olig1/Olig2* as 0.89 ≤ *γ* ≤ 1.42. This approach was used to identify the range of the structural resolution parameter for each 5C region for each cell type.

We next sought to comprehensively identify the possible communities across length scales within our 5C regions. We ran the Louvain-like algorithm and consensus similarity method across our computed *γ* ranges in increments of 0.01. As predicted, we detected smaller communities at higher *γ* values nested within larger communities detected at a lower *γ* values (**Figure 3D**). Many communities corresponded with domain structures readily observable in the heatmaps, however there were domains identified that qualitatively did not correspond with domain structures. These data suggest that the modularity optimization and consensus similarity approaches can successfully identify a nested hierarchy of community structures and highlight the need of additional methods to remove low-confidence communities from the full detected set.

### Refinement of community calls via hierarchical variance minimization

Upon closer examination of the partition blocks across values of the structural resolution parameter, we noticed that the community calls that aligned more closely with domain structures in the 5C heatmaps exhibited largely consistent community boundaries across the 100 partitions (**Supplementary Fig. 3A+B**). By contrast, community calls that did not align with domains tended to exhibit greater variability in boundary assignment across the 100 partitions (**Supplementary Fig. 3A+C**). On the basis of these observations, we posited that degeneracies in community assignments across 100 runs of the modularity optimization algorithm could point to a low probability community call, potentially indicative of overlapping or soft community structure (Ball, Karrer, and Newman 2011). To test this hypothesis, we devised a method in which we preferentially selected robust communities with low boundary variance from among all the partitions (**Supplementary Fig. 4**). We first set out to identify partitions that were stable across multiple resolution parameter values. By plotting the average number of communities versus the resolution parameter (Bassett et al. 2013), we uncovered plateaus in which adjacent *γ* values gave rise to the same average number and spatial placement of communities. (**Figure 4A, B**). Additionally, we noticed that some *γ* values did not belong to a plateau, suggesting that the underlying community structures predicted by these partitions are not robustly detected across resolution parameter values and may represent weak community boundaries. Thus, we implemented a strategy in which we pooled the partition blocks with the same average number of communities into a single ‘hierarchy level’ and then calculated a consensus partition at each level (**Figure 4C**). Partition blocks not in plateaus were excluded from downstream analyses.

**Figure 4.**
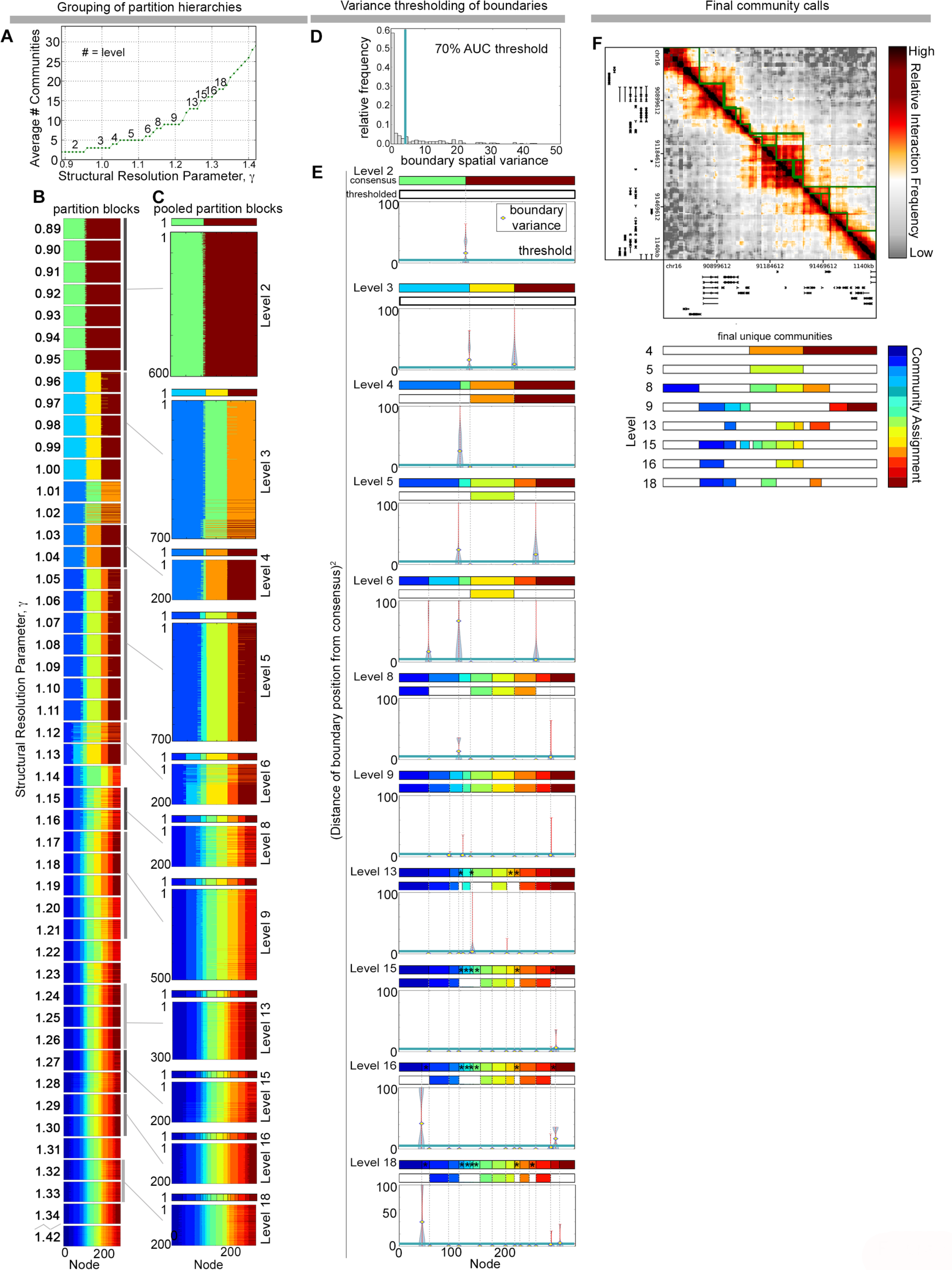
Hierarchical spatial variance minimization method identifies *bona fide* communities within partitions. (A) Average number of communities across 100 partitions *versus* resolution parameter for the *Olig1-Olig2* network. Groups of two or more consecutive *γ* values with the same average number of communities are grouped into levels. (B) Sweep of partition blocks across the range of *γ* (left). (C) Two or more consecutive partition blocks from (B) that have the same average number of communities are grouped into levels. The consensus partition of each pooled partition block is shown above its corresponding pooled partition block. (D) Distribution of boundary spatial variance pooled across all levels and all 5C genomic regions for pNPC cells. The chosen boundary variance threshold of 4.19, which corresponds to 70% Area Under the Curve (AUC) is shown in blue. (E) Thresholding of communities based on the 70% AUC boundary variance threshold. For each level, consensus partitions prior to thresholding are shown, beneath which are variance thresholded consensus partitions. Black asterisks over communities in the consensus partitions prior to thresholding indicate that the community is ≤ 48*kb* and does not contribute to the global boundary variance distribution in panel (D). These communities are removed from further consideration. Violin plots of the squared differences per boundary, 
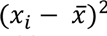
, are shown for each level, where *x_i_* is the boundary coordinate for a given partition amongst the pooled partition block and 
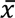
 is the boundary coordinate of the consensus partition. Boundary variance is indicated by a yellow diamond. The boundary variance threshold of 4.19 is drawn as a horizontal blue line on the violin plots. Note that the x-axis coordinates are the same for the consensus plots and the violin plots; light grey dashed lines connect a given consensus boundary to the violin plot of the squared differences for that boundary. A community that has at least one of its two boundaries with a variance greater than the threshold are removed from the list of final community calls. (F) Final unique community calls of variance-thresholded consensus partitions.

Using the genomic location of the consensus communities as the expectation, we next computed a spatial variance for each community boundary within each consensus partition at each level. To ensure our threshold was robust to hierarchy level and genomic region, we removed communities that were smaller than 48 kb in size and pooled the boundary variances across hierarchy levels and across the seven ~1 Mb genomic regions covered in our 5C data set. We observed that the boundary spatial variance distribution was zero-inflated, indicating a high number of consistent community boundaries across applications of the Louvain-like locally greedy algorithm to maximize network modularity (**Figure 4D**). We then removed communities with at least one boundary exhibiting a spatial variance greater than 4.19, the value corresponding to the 70% AUC (**Figures 4D, E**). Lastly, we removed redundant communities with identical boundaries. The resulting final partition is consistent across multiple structural resolution parameters and multiple applications of the Louvain-like locally greedy algorithm. Thus, our new method of ‘hierarchical spatial variance minimization’ results in the elimination of low probability community boundaries to focus on final communities that align well with the subTAD domain structures in the heatmaps (**Figure 4E**).

### Sub-Mb scale communities are markedly different across genomic regions and cell types

We next sought to determine if our hierarchical variance minimization method was robust to genomic loci with markedly different architectural features. We applied our method to all ~1 Mb genomic regions in our 5C library in pNPCs and ES cells. The resulting number, placement, and size of the hierarchical community structures were markedly different across the genomic regions queried **(Figure 5A-G)**. Spring force diagram visualizations of the genome topology networks further highlight the differences in topological organization and community assignments across the three genomic regions **(Figure 5 H-M)**. Community sizes ranged from 52 to 684 kb with median sizes in *Olig1-Olig2*, *Sox2*, and *Klf4* regions of 96, 96, and 84 kb for pNPCs and 108, 112, 104 for ES cells, respectively (**Figure 5 N-O)**. These results highlight that our community detection method can identify communities in a manner that is robust to a range of disparate architectural features and does not require *a priori* knowledge of the number or size of communities.

**Figure 5.**
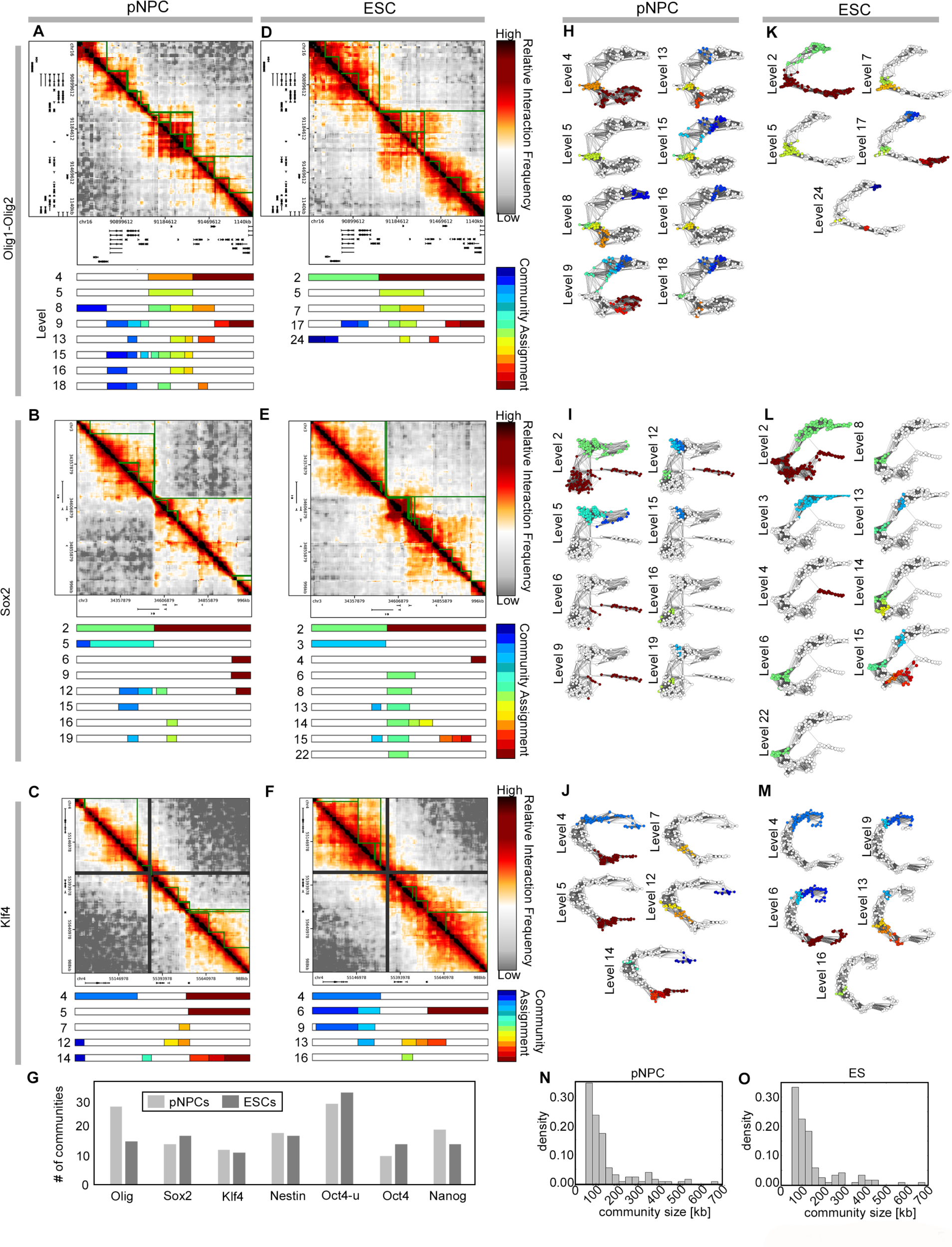
Hierarchical spatial variance minimization captures folding differences across genomic loci and two cellular states. (A-F) Final community calls using hierarchical variance minimization for ~1Mb regions surrounding Olig1-Olig2, Sox2, and Klf4 in primary neural stem cells (pNPCs) and embryonic stem cells (ESCs). (G) The number of communities identified in each of the seven 5C genomic regions across the two cellular states. (H-M) Spring force diagrams of the Klf4, Sox2, and Olig1-Olig2 networks across levels for pNPCs and ESCs. The colors of the nodes indicate community assignment of the node and correspond to the colors in the thresholded consensus partitions shown in panels (A-F). (N-O) Histograms of community sizes identified in pNPCs and ESCs, respectively, across the seven 5C genomic regions.

We then set out to identify the communities that exhibit statistically significant changes between ES cells and pNPCs. Qualitatively, we observed that there were marked differences in the communities identified in the two cell types **(Figure 5A-F)**. We next constructed our statistical test with a null hypothesis that a given community of interest is constitutive and thus the aRAND computed between the community in ES cells and pNPCs is unity. We used two replicate 5C data sets for each of our cell types to construct a set of hybrid pseudo-replicates (see **Materials and Methods**). For a given cell type, we then sampled the pseudo-replicates 10,000 times with replacement and computed the aRAND between a given community of interest in our replicate of interest (ES or pNPC) and the most similar community identified in the hybrid pseudo-replicate. This bootstrapped sampling procedure resulted in a distribution of similarity scores for each community under the null hypothesis that the community is constitutive. The position of the aRAND index computed between pNPC and ES cells for the particular community in question was compared to its bootstrapped null distribution. (For a detailed description, see **Materials and Methods**). For *p*-values < 0.05, we identified 45 ES-specific and 66 pNPC-specific communities for which we could reject the null hypothesis (**Figure 6A, B**).

**Figure 6.**
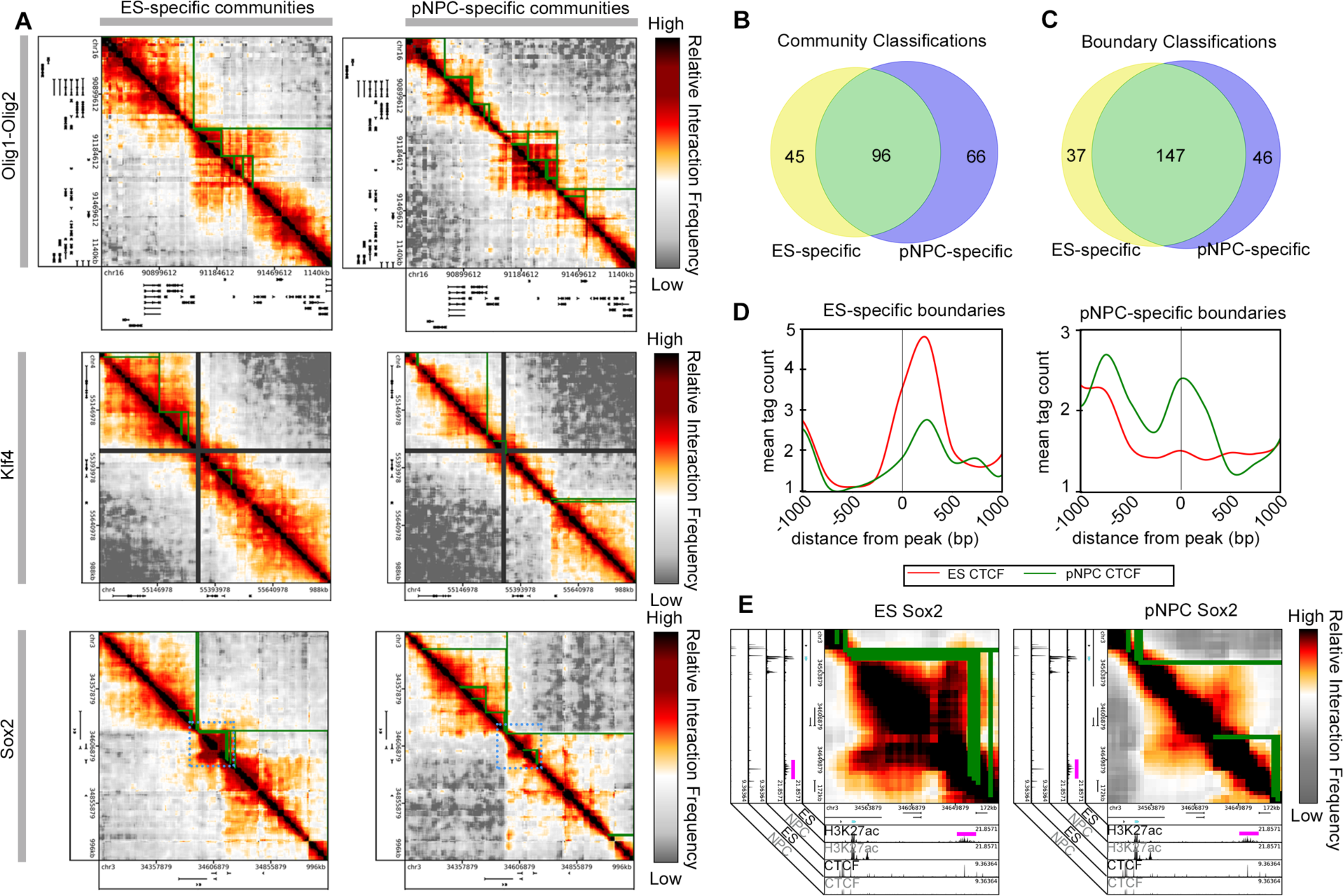
Statistical testing identifies dynamic communities across two cellular states. (A) Significantly dynamic communities identified across two cellular states (embryonic stem and primary neural progenitor) are drawn in green on 5C heatmaps for three genomic regions. Dashed blue boxes on the Sox2 heatmaps indicate the zoom-in region shown in panel (E). (B, C) Venn diagram of panel (B) community and panel (C) boundary classifications across all seven 5C genomic regions. (D) Average CTCF ChIP-seq signal at cell-type specific community boundaries. (E) Close-up view of community structure surrounding the *Sox2* gene, which is colored in blue. H3K27ac and CTCF ChIP-seq tracks from ES cells and pNPCs are shown beneath the heatmaps. The ES-specific super-enhancer is indicated with a pink bar above the ES H3K27ac ChIP-seq track.

To further confirm the biological relevance of our communities, we identified an ES-specific community containing the key pluripotency gene, *Sox2*, and its ES-specific super-enhancer that is known to play a critical role in maintaining the pluripotency transcription program in ES cells (**Figure 6E)** (Li et al. 2014). Neither the super-enhancer nor the community bounded by the super-enhancer and *Sox2* is present in neural progenitor cells (**Figure 6E)**, thus high-lighting the biological relevance of the dynamic communities we identified. Moreover, among these cell type specific communities, we identified dynamic community boundaries and found 37 ES-specific and 46 pNPC-specific boundaries (**Figure 6C**). ES-specific boundaries were enriched for ES-specific ChIPseq signal for the architectural protein CTCF, whereas pNPC-specific boundaries were enriched for pNPC-specific CTCF (**Figure 6D**). Overall, by combining the Louvain-like locally greedy algorithm for modularity maximization with our newly developed hierarchical spatial variance minimization method and a bootstrapped statistical test, we detect subMb-scale, nested domains within TADs, some of which are markedly altered between ES cells and pNPCs and correlate with known switches in gene expression regulation.

## Discussion

Complex systems from a wide range of orthogonal disciplines can be represented as networks. Networks are defined mathematically as a collection of nodes and edges between nodes. Because 3-D genome topology is queried by proximity ligation assays that measure the interaction frequency (edge weight) between genomic loci (nodes), we reasoned that tools from network mathematics might have utility in capturing the conformation of higher-order chromatin structure. By applying a modularity optimization algorithm to HiC and 5C data, we find that the topology of the mammalian genome is organized into hierarchically nested community structure at the subMb scale within TADs. To date, the precise quantification of the nested domains within TADs has been limited by the accuracy and/or sensitivity of the metrics employed and confounding effects from the larger, higher-order TAD and compartment folding patterns. Here, we demonstrate that larger TADs and smaller subTADS can be parsed across length scales via the fine-tuning of the resolution parameter in a Louvain-like locally greedy modularity maximization algorithm. In addition to developing a new method for sensitively detecting subTADs, we also applied this method to quantify dynamic alterations in subTAD community structures between two developmentally relevant cellular states. We found that our method could capture cell-type specific switches in community organization. Notably, we identified cell-type specific communities containing developmentally regulated genes and cell-type specific enhancers in pluripotent ES cells that switch community organization upon differentiation to pNPCs, a feature reminiscent of the modular miRNA co-expression profiles during reprogramming (Henzler et al. 2013). Domain boundaries mirror occupancy patterns of a known architectural protein, lending independent support to the validity of our community detection method. Together, these data suggest that genome folding can be abstracted as a network and that graph theory methods have utility in capturing some of the more nuanced features of fine-scale genome folding within TADs.

The mammalian brain is an example where network science has been applied to yield key insights into the relationship between structure and function in biology (D. S. Bassett and Bullmore 2006; E. T. Bullmore and Bassett 2011). In large-scale modeling of neuronal connectivity in the brain, a node typically represents a subregion of the brain. Edges between nodes could represent anatomical (e.g. density of white matter tracts between brain regions) or functional (e.g. temporal correlations in activity) connections. A critical insight obtained by the application of graph theory to the brain has been the identification of large-scale structures termed modules, or communities (Sporns and Betzel 2016). These modules are believed to aid in specialized or segregated computations (Bullmore and Sporns 2012) in complex information processing systems (Simon 1962). In the context of brain anatomy, a module is a subnetwork of neurons across multiple tissues that exhibit dense connections compared to the larger network, and these subnetworks can be composed of smaller and smaller subnetworks in a hierarchically nested architecture similar to that we observe here (Danielle S. Bassett et al. 2010; Danielle S. Bassett, Brown, et al. 2011). Similarly, even if brain regions exhibit sparse anatomical connections, they might display highly correlated temporal patterns in functionality, indicating a densely connected functional community structure that also displays hierarchical nesting (Salvador et al. 2005; Meunier, Lambiotte, and Bullmore 2010). Functional and structural communities have been found to play a role in important biological and cognitive processes such as learning (Bassett et al. 2011; Bassett et al. 2013) and aging (Meunier et al. 2009; Gu et al. 2015).

Here, in a cell type that serves as a precursor to many mature cell types in the brain, we zoom in to the nucleus of cells that make up the larger scale brain networks, and we model the network properties of the genetic material. Our observation that one of the salient features of brain network organization – hierarchical modular architecture – is mirrored in the genetic material of neuronal cell types, motivates the question of what evolutionary benefit modularity offers to both systems. Prior work in evolution and development of other biological systems has suggested that modular architecture supports evolvability of a system, because one portion (or module) can evolve relatively independently of other modules, thereby minimizing adverse effects on system function (Kirschner and Gerhart 1998). Indeed, because modularity supports evolvability, it also likely originates in the first place as a natural response to the network’s need to satisfy rapidly changing goals (Kashtan and Alon 2005) and to generalize its learned responses to new environments (Parter, Kashtan, and Alon 2008) through the theory of facilitated variation (Gerhart and Kirschner 2007). It will be interesting in the future to determine how the modules identified here might relate to the modular nature of gene expression, which changes appreciably over evolutionary time scales (Conaco et al. 2012).

An important next step is to conduct causal genetic studies to determine the functional purpose of our identified genome folding communities. In tandem with these empirical studies, it will be important to develop quantitative theories of the role of folding patterns on system function. Particularly promising avenues include Boolean modeling (Wang, Saadatpour, and Albert 2012), which offers a qualitative and phenomenological framework to infer biological regulatory mechanisms in biological systems wherein the knowledge of mechanistic details and kinetic parameters is scarce. Furthermore, it will be interesting to use generative modeling approaches (Betzel et al. 2016) to identify the topological pressures that explain the most variance in the observed hierarchically modular configurations. The long-term goal for these efforts would be to create theoretical models that bridge disconnected length scales to understand the potential causative link between genome structure-function and more global brain structure-function (Arcila et al. 2014).

In this work, we describe a new approach in which we apply graph theory to accurately capture fine-scale folding patterns of the mammalian genome. Notably, we quantify the dynamic reconfiguration of fine-scale topological domains as embryonic stem cells switch cellular fate to neural progenitor cells. These results are significant because it is widely thought that fine-scale genome organization within TADs plays a critical role in gene expression changes across development. Although large scale genome folding domains (so-called TADs) have been accurately quantified, there is still no robust, gold-standard method for quantifying the nested properties of chromatin domains at the sub-Mb scale within TADs. The ability to accurately quantify nested, hierarchical, fine-scale genome organization and dynamics may shed new light into how genome folding governs genome function across development and during the onset and progression of disease.

## Code availability

The code necessary to run the method as well as full usage instructions will be publicly available on Bitbucket upon acceptance to a peer-reviewed journal.

## Materials and Methods

### Hi-C mapping, normalization, and binning

Hi-C data from the mouse cortex generated in Dixon et al. 2012 was downloaded from GEO from accession number GSE35156 (Dixon et al. 2012). Paired-end reads were aligned independently to mm9 mouse genome using bowtie2 (global parameters: ‐‐very-sensitive ‐L 30 ‐score-min L,−0.6,−0.2 ‐end-to-end ‐‐reorder; local parameters: ‐‐very-sensitive ‐L 20 ‐score-min L,−0.6,−0.2 ‐end-to-end ‐‐reorder) through the HiC-Pro software (Servant et al., 2015). Unmapped reads, non-uniquely mapped reads and PCR duplicates were filtered and uniquely aligned reads were paired. HiC maps were generated at 40 kb matrix resolution and balanced using the iterative correction and eigenvector decomposition (ICED) technique (Imakaev et al., 2012). The resulting genome-wide Hi-C data is represented in a matrix format *A_ij_*, where the *ijth* element represents the interaction frequency between 40 kb bin *i* and bin *j.*

### Hi-C data network construction

Mapped, normalized Hi-C data were parsed into 60 Mb regions surrounding *Sox2* and *Olig1-Olig2* genes key to the neural progenitor cell state. The resulting contact matrices represent weighted, undirected networks where each bin represents a node and each bin entry of ‘Relative Interaction Frequency’ represents a weighted edge.

### 5C mapping, normalization, and binning

5C libraries generated in Beagan et al. in pluripotent mouse ES cells and multipotent neural progenitor cells were downloaded from GEO accession numbers GSM1974095, GSM1974096, GSM1974099, and GSM1974100 (Beagan et al. 2016). Data were processed as described in detail in the Supplementary Methods of Beagan et al. (Beagan et al. 2016). Briefly, paired-end reads were mapped to a pseudo-genome consisting of all 5C primers using Bowtie (<u>http://bowtie-bio.sourceforge.net/index.shtml</u>) (Langmead 2010). Reads with redundant alignments were removed and interactions between forward and reverse primer pairs were counted as the number of unique counts that map to the pseudo-genome on both ends of the paired-end read. Counts were converted to contact matrices for each genomic region queried (i.e. ~1 Mb regions around developmentally regulated genes *Sox2* (chr 3: 34108879 – 35104879), *Olig1-Olig2* (chr 16: 90614612 – 91754612), *Klf4* (chr 4: 54899978 – 55887978), *Nanog* (chr 6: 122189359 – 122737359)*, Nestin* (chr 3: 87285087 – 88369087), *Oct4-upstream* (chr 17: 34430391 – 35474391) and *Oct4-downstream* (chr 17: 35630391 - 36062391).

To correct for biases in the restriction fragments and batch effects, we quantile normalized and primer corrected the data as previously described (Beagan et al. 2016). Data were then binned into 4 kb bins and smoothed by replacing the counts within each 4 kb with the average count within a 24 kb window centered at that bin. Binned and smoothed 5C data is referred to as “Relative Interaction Frequency”. Primers that contained less than 100 counts across all potential primer pairs were converted to NaN. Similarly, individual primer-primer pairs with less than 9 counts post primer normalization were converted to NaN. NaNs were subsequently smoothed by substituting NaN values with the value of the mean of the 8 x 8 surrounding pixels (or 16 kb).

### 5C data network construction

Similar to Hi-C data, 5C contact matrices represent weighted, undirected networks where each bin represents a node and each bin entry of ‘Relative Interaction Frequency’ represents a weighted edge.

### Community detection: Network Modularity Maximization using Louvain-like Algorithm

To partition our genome folding networks into communities, we employed a Louvain-like, locally greedy algorithm to maximize modularity. We first computed the modularity matrix, *M*, according to (**1**)

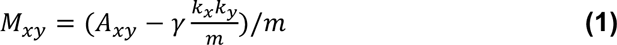

where *M* is a *CxC* matrix, *C* is the total number of communities, *M_xy_*,[ is the normalized interaction frequency between *x* and *y* communities, *k_x_* is the sum of all edge weights for community *x*, *γ* is a structural resolution parameter (discussed below) and *m* is the total sum of all edge weights in network *A* (i.e., the sum of all interaction counts in the matrix). For the first iteration of the algorithm (discussed in detail below), each individual node is an independent community, thus indices for communities *x* and *y* correspond the indices for nodes *i* and *j* (see **Equation 1, Results**). For any given resolution parameter value, *A_xy_*/*m* is the normalized strength of edges that connect communities *x* and *y*, and 
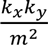
 is the expected normalized strength of edges at communities *x* and *y* if edges are placed at random. From the modularity matrix, *M*, the modularity, *Q*, is computed according to (**2**):

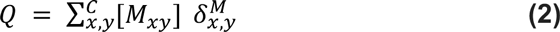

where *M_xy_*, is the interaction frequency between *x* and *y* communities normalized to the expected interaction probability, *C* is the total number of communities in matrix *M* at each iteration of the algorithm (discussed below). To ensure that only edges within communities are added to the summation, 
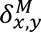
 is 1 if *x* and *y* are assigned to the same community and 0 otherwise.

The assumption of the Louvain-like algorithm is that the optimal community structure can be resolved by maximizing *Q*. The algorithm relies on an iterative, dynamic programming approach to rapidly converge on a local maximum in *Q* without comprehensively examining the entire search space. At each iteration, *t*, each individual node is given the opportunity to move into a new community placement that yields a locally maximal aggregate gain in *Q*, *ΔQ*, for the *M* network according to (**3**):

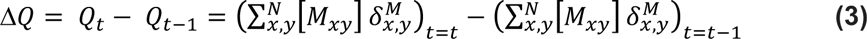

In the first iteration, *t* = 0, matrix *M* has the same dimensions as matrix *A* such that every node is assigned to its own community (i.e. *C* = *N* and *x* = *i*). Each node is given the chance to merge and at the end of the iteration, if *ΔQ* ≤ 10^−10^, the algorithm terminates on the basis of the assumption that no further moves will lead to a notable increase in *ΔQ*. If *ΔQ* > 10^−10^, then the algorithm proceeds to the next iteration and resizes the modularity matrix to *CxC* dimensions (where *C* is the number of communities computed in the previous *t* = *t* − 1 iteration). The intuition for the iterations is that with increasing *t* the average size of communities increases with the gain in modularity. The algorithm terminates when no further single community merge leads to an improvement in modularity. Due to the tendency of modularity maximization algorithms to converge on local maxima (Good, de Montjoye, and Clauset 2010), the Louvain-like algorithm was applied 100 times to sample the landscape of local maxima for each resolution parameter value. To ensure random sampling of the local maxima landscape, each of the 100 partitions is run with a randomized node selection order with unique seed value.

### Computing the aRAND index as a similarity metric for partitions

To compare the similarity between any two given community partitions of a network, we used an adjusted RAND index (aRAND) metric from the sklearn toolbox (sklearn.metrics.adjusted_rand_score, Traud et al. 2008; Hubert and Arabie 1985). This metric has utility over the classic RAND index because it accounts for the similarity between two community partitions expected by random chance (Bassett et al. 2013). For any two community partitions, *X* and *Y*, the aRAND is computed according to (**4**):

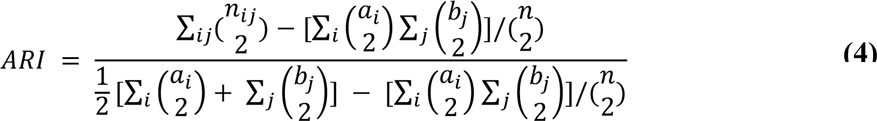

where *n_ij_* is the number of nodes in common between community *i* in partition *X* and community *j* in partition *Y*, *a_i_* is the number of nodes in community *i*, *b_j_* is the number of nodes in community *j*, and *n* is the total number of nodes in the network. Equation (**4**) can be represented as (**5**):

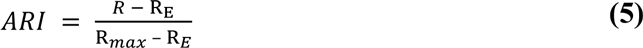

where *R* is the number of outcomes in which node pairings are in the same community in *X* and in the same community in *Y*, *R_max_* is the number of outcomes a node pairing can be placed in the same community for a given partition, and *R_E_* is the background correction factor, or the random chance that a pairing in the same community will also occur in the same community for another partition. For perfectly matched partitions, *ARI* is equal to unity. For partitions that are no more similar than would be expected by random chance, the ARI is close to 0 or slightly negative.

### Determination of a consensus partition

To identify a consensus partition among a set of partitions, we computed the aRand between all pairs of partitions within the set. The partition with the highest average aRand, representing the partition that is most similar to all other partitions, was selected as the consensus partition (Doron, Bassett, and Gazzaniga 2012; Lohse et al. 2014).

### Boundary spatial variance calculation

To quantify the variability of a community boundary call across partitions within a set of partitions, we defined a novel approach based on boundary spatial variance minimization. Specifically, we computed the spatial variance per boundary across the 100 partitions at a given resolution value according to (**6**):

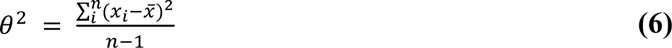

where *x_i_* is the coordinate of the boundary in partition *i* (in units of nodes), 
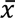
 is the coordinate of the consensus boundary, and *n* is the number of partitions in the set. Boundaries with perfect agreement across the set of partitions will have a variance of zero, whereas boundaries with large fluctuations in position across the set of partitions will have a higher variance.

### *Community detection of Hi-C data* and comparison to gold-standard TADs

We applied the Louvain-like locally greedy algorithm for modularity maximization at a range of resolution parameter values to detect communities of different sizes in mouse cortex Hi-C. We identified a consensus partition (computed as described above) from the 100 partitions resulting from running the modularity optimization algorithm 100 times at a given resolution value. To quantify the performance of our approach in the identification of TADs, we constructed a receiver operating characteristic (ROC) curve by ranking community boundaries identified at *γ* = 4.1, the resolution value at which community calls most closely mirrored the gold standard TAD calls, across the two 60 Mb regions by their spatial variance and comparing them to gold standard TAD calls (Dixon et al. 2012). We computed the area under the ROC curve as a metric of performance that reflects both the sensitivity and specificity of our TAD calls.

### Computing region-specific range of the resolution parameter

To compute the range of the resolution parameter for each 5C region, we first constructed a randomly rewired network by reshuffling the edge weights in a manner that preserves the symmetric nature of the adjacency matrix, as well as preserving node degree while approximately preserving node strength (Rubinov and Sporns 2011; Maslov and Sneppen 2002). To create this randomly rewired network, we used null_model_und_sign.m tool from the Brain Connectivity Toolbox (https://sites.google.com/site/bctnet/). Briefly, the expected weight, *E_i,j_*, for all edges is computed as *k_i_* * *k_j_* where *k_i_* is the degree of node *i* and *k_j_* is the degree of node *j*. Observed (*A_i,j_*) and expected (*E_i,j_*) edge weights are then rank ordered in two independent lists. We then proceed with the following randomization steps: (1) A specific edge index *i*, *j* (where *i* ≠ *j*) is randomly selected, (2) the rank of that edge in the expected list is noted, (3) the observed edge weight at the noted expected rank is then assigned to edge *A_i,j_*. (4). The expected and observed values are then removed from their respective lists. A new set of expected values is computed and then ranked to reflect the removed edge. (5) Steps 1 – 4 are repeated until all observed edge values have been assigned. Using the above algorithm, we constructed 10 randomly rewired networks for both cellular conditions (i.e. pNPC and ES cells) across all 7 genomic regions queried by 5C.

A range of resolution parameter, *γ*, values was explored for both real and randomly rewired networks. For each *γ* value in the range of 0.8 to 1.6 in increments of 0.01, modularity *Q* was computed for one run of the Louvain algorithm for the real and randomly rewired 5C networks. We identified our range of gamma as the *Q* where the real network diverges and converges from the random network, as assessed by the *γ* values where *Q* of the real network is greater than 15% of the maximum difference between *Q_real* and *Q_random*.

### Hierarchical grouping and spatial variance thresholding for refined community calls

To identify high confidence communities, we developed a ‘hierarchical spatial variance minimization method’. Initially, we performed community detection across the range of resolution values for a given region, resulting in a ‘partition block’ or set of 100 community partitions for each resolution parameter value. For a given region, we grouped partition blocks with the same average number of communities together to form a ‘hierarchy level’ and then determined a consensus partition for each hierarchy level. We next computed the spatial variance at the boundaries of all communities > 48 kb (a community size threshold we set because it corresponds to 2x our smoothing window) and pooled the values across levels and regions for a given cell type. We used the distribution of boundary variance values across all of our seven 5C regions to select a variance threshold that corresponded to 70% Area Under the Curve (AUC). Communities with one or more boundaries that did not pass the variance threshold were considered low confidence and were removed from the list of community calls. Because communities with the same start and stop coordinates may be called more than once across hierarchies within a region, we removed non-unique community calls (identical start and stop coordinates). The resulting unique communities represent the high confidence final list of communities called for a given cell type.

### Community-specific aRAND

To compute the similarity between two communities, *x* and *y*, we first converted community partitions into single-community binary partitions for a given community of interest by creating an array, *C*, with the number of elements equal to the number of nodes in the network. If node *i* belongs to the community of interest, *C_i_* = 1, whereas if node *i* does not belong to the community of interest, *C_i_* = 0. The similarity between two communities is then determined by calculating the aRAND on the two binary community partitions, *C_x_* and *C_y_* (Traud et al. 2008; Hubert and Arabie 1985).

### Community classification across cell types

To identify communities that reconfigure across cellular states in a statistically significant manner, we first constructed 200 pseudo replicates for each cell type by randomly hybridizing between the two available 5C replicates, *A* and *B*, for each cell type. We constructed a given pseudo replicate hybrid network, *H*, by randomly determining whether edge *H_ij_* in the upper triangular matrix is assigned the value *A_ij_* or *B_ij_* and then mirroring the edge weights of *H* across the diagonal. We next identified high confidence communities for each of the 200 pseudo replicates of each cell type by applying the network modularity maximization followed by hierarchical variance minimization method. For each high confidence community identified in the real replicate, *A*, we constructed a bootstrapped, community-specific null distribution sampling the 200 pseudo-replicates 10,000 times with replacement. For each sampling, we identified the pseudo-replicate community most similar to the community of interest by first ranking the pseudo-replicate communities by the number of nodes they share in common with the community of interest, then selecting the top 50% of ranked pseudo-replicate communities, and lastly computing the community-specific aRAND between the community of interest and the selected pseudo-replicate communities (if any). The maximal community-specific aRAND corresponds to the most similar community (if no overlapping community is identified the aRAND is zero), and the aRAND is appended to the null distribution.

We next evaluated the cell-type specificity of a community of interest by identifying the most similar community in the other cell type, computing the community-specific aRAND, and assessing the position of the aRAND in the null distribution to compute a *p*-value according to **(7)**:

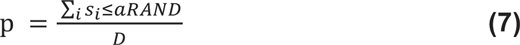

where *D* is the number of values in the null distribution and *s_i_* is a value in the null distribution. We considered communities with a *p*-value < 0.05 to be significantly dynamic across the cellular states.

### Identification of dynamic boundaries

To identify a list of dynamic community boundaries, we filtered the list of boundaries of dynamic communities by removing boundaries that were present in both cell types. This was necessary because some boundaries may be present in both cell types but the communities they border may be considered dynamic.

### Spring force diagram visualization

We visualized networks as spring-force diagrams using the Kamada Kawai spring force algorithm (Fruchterman and Reingold, Code from MatlabBGL toolbox, David Gleich). For visualization purposes, we threshold the networks such that the top 15% of edge weights remained. A threshold of 15% was chosen because it was stringent enough to improve visualization, but it was lenient enough for the graph to remain fully connected. Note that all analyses were performed on the fully weighted graphs; thresholds were applied only for visualization and not for analysis.

## Acknowledgements

J.E.P.C. is a New York Stem Cell Foundation (NYSCF) Robertson Investigator and an Alfred P. Sloan Foundation Fellow. This work was funded by The New York Stem Cell Foundation (J.E.P.C), the Alfred P. Sloan Foundation (J.E.P.C, D.S.B), the NIH Director’s New Innovator Award from the National Institute of Mental Health (1DP2MH11024701; J.E.P.C), a 4D Nucleome Common Fund grant (1U01HL12999801; J.E.P.C), and a joint NSF-NIGMS grant to support research at the interface of the biological and mathematical sciences (1562665; J.E.P.C). This material is based upon work supported by the National Science Foundation Graduate Research Fellowship under DGE-1321851 (H.K.N). D.S.B. would also like to acknowledge support from the John D. and Catherine T. MacArthur Foundation, the National Institute of Child Health and Human Development (1R01HD086888-01) and the National Science Foundation (BCS-1441502, PHY-1554488, and BCS-1631550).

## Supplementary Figures

**Supplementary Figure 1.**
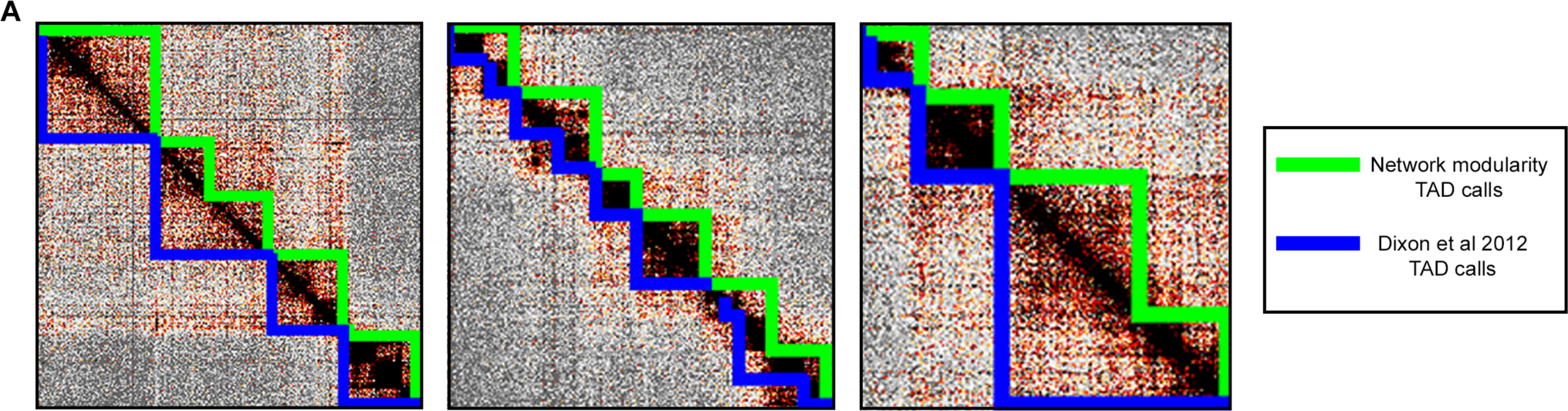
Dixon et al. TAD calls and Network Modularity TAD calls form nested calls at the sites of disagreement. (A) TAD calls from Dixon et al. 2012 are outlined in blue. TAD calls using Network Modularity maximization are outlined in green.

**Supplementary Figure 2.**
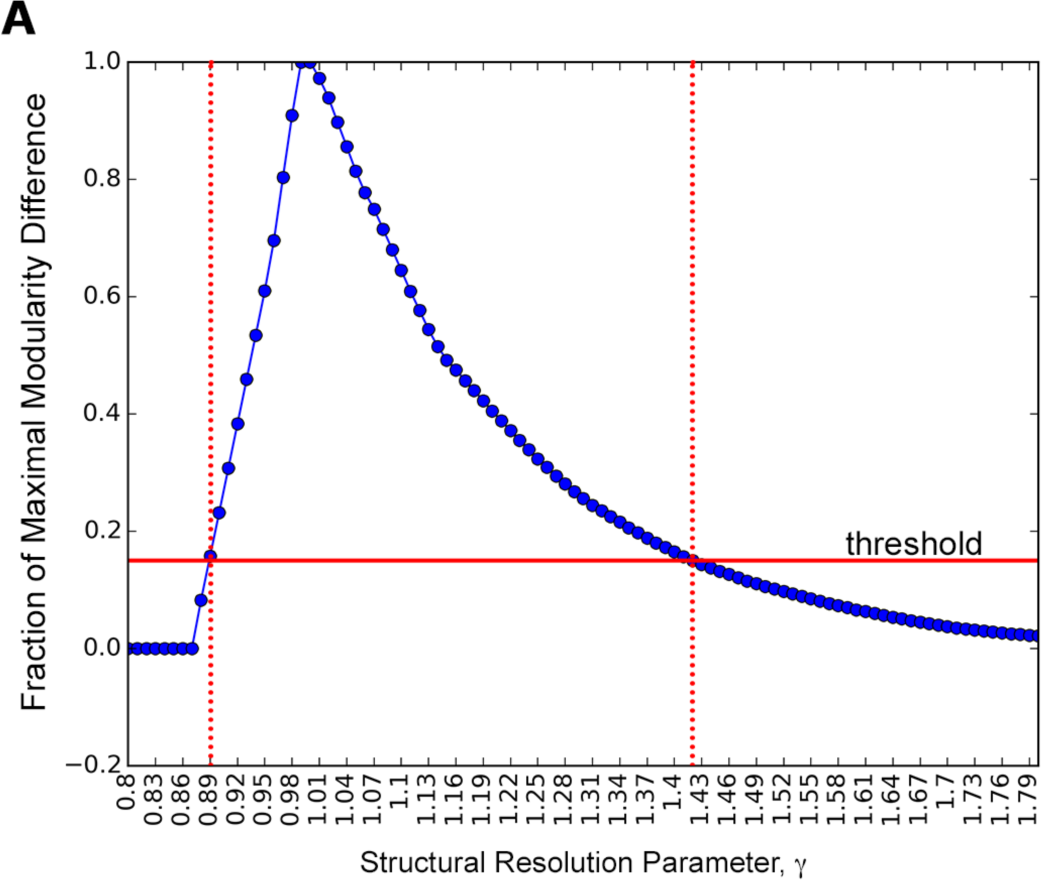
Determination of dynamic range of structural resolution parameter, *γ*. The difference in modularity, *Q*, values between the pNPC Olig1-Olig2 network and 10 randomly re-wired networks normalized to the maximal difference. The red horizontal line indicates 15% of the maximal difference in *Q* between real and random networks. Structural resolution parameter, *γ*, values between the two vertical red dashed lines are used for modularity maximization.

**Supplementary Figure 3.**
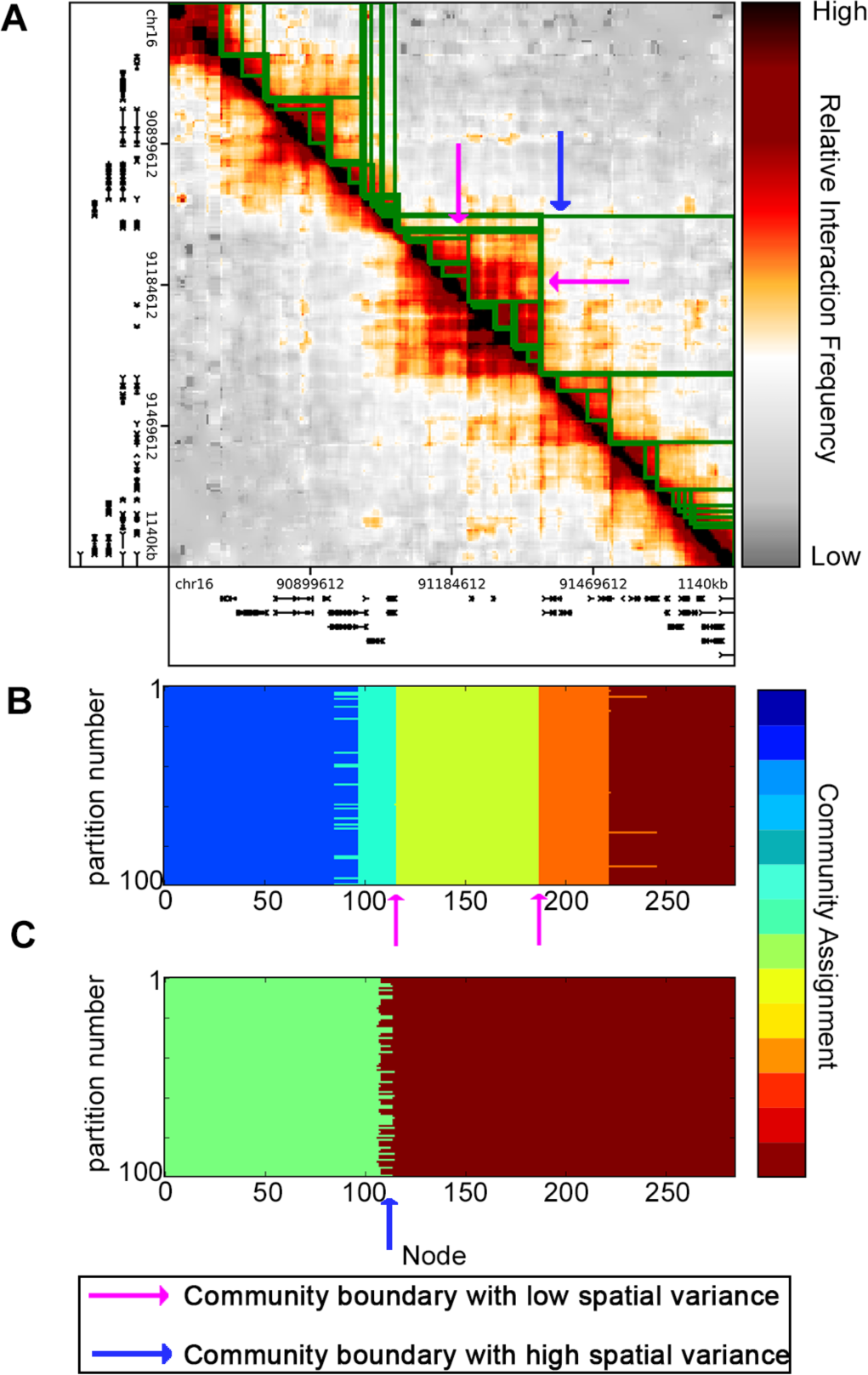
Partition blocks with high and low variability in boundary position. (A) Input pNPC Olig1-Olig2 network data represented as a matrix of relative interaction frequencies between nodes. (B) One hundred community partitions at the same structural resolution parameter, *γ*, value. Boundaries with low spatial variance are indicated by the pink arrows. (C) One hundred partitions at a structural resolution parameter, *γ*, value different from that of panel (B). A boundary with high spatial variance is indicated by the blue arrow.

**Supplementary Figure 4.**
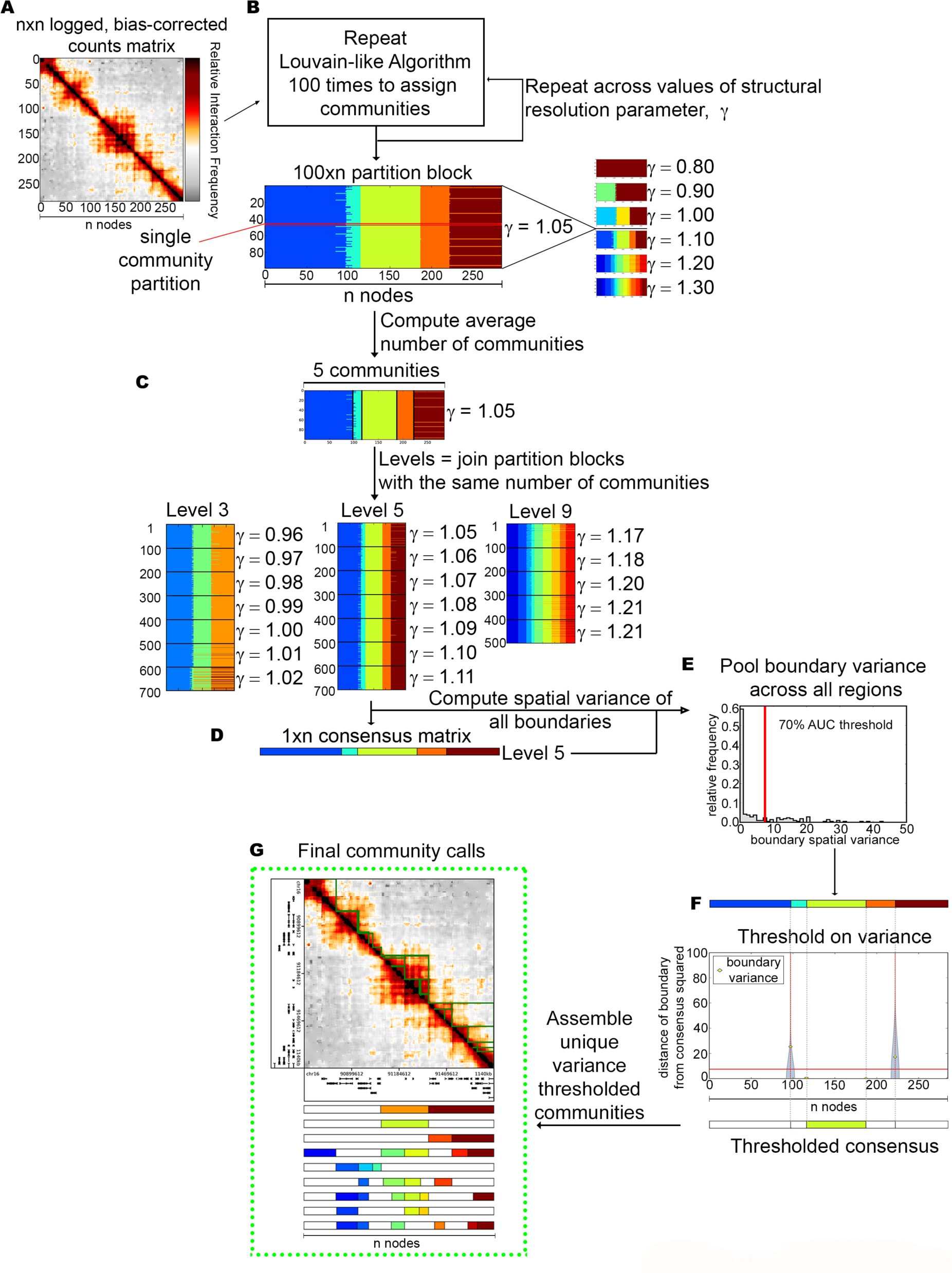
Overview of Hierarchical Spatial Variance minimization method. (A) Input data is an n x n matrix of relative interaction frequencies between genomic loci, where n is the number of nodes. (B) The Louvain-like greedy algorithm is repeated 100 times for a range of structural resolution parameter, *γ*, values. (C) Partition blocks with the same average number of communities are grouped into hierarchy levels. (D) A similarity consensus partition is chosen for each hierarchical level. (E) The boundary spatial variance is computed for each boundary at each level for all regions under investigation for a given cell type. A boundary spatial variance threshold corresponding to 70% Area under the Curve of the distribution of pooled boundary spatial variance values is selected. (F) Communities whose boundaries do not pass the variance threshold are removed from the list of communities. (G) The final community calls consist of unique communities > 48 kb that have passed the variance threshold.

